# Genome-wide characterization of T cell responses to *Bordetella pertussis* reveals broad reactivity and similar polarization irrespective of childhood vaccination profiles

**DOI:** 10.1101/2023.03.24.534182

**Authors:** Ricardo da Silva Antunes, Emily Garrigan, Lorenzo G Quiambao, Sandeep Kumar Dhanda, Daniel Marrama, Luise Westernberg, Eric Wang, Aaron Sutherland, Sandra K Armstrong, Timothy J Brickman, John Sidney, April Frazier, Tod Merkel, Bjoern Peters, Alessandro Sette

## Abstract

The incidence of whooping cough (pertussis), the respiratory disease caused by *Bordetella pertussis* (BP) has increased in recent years, and it is suspected that the switch from whole-cell pertussis (wP) to acellular pertussis (aP) vaccines may be a contributing factor to the rise in morbidity. While a growing body of evidence indicates that T cells play a role in the control and prevention of symptomatic disease, nearly all data on human BP-specific T cells is related to the four antigens contained in the aP vaccines, and data detailing T cell responses to additional non-aP antigens, are lacking. Here, we derived a full-genome map of human BP-specific CD4+ T cell responses using a high-throughput *ex vivo* Activation Induced Marker (AIM) assay, to screen a peptide library spanning over 3000 different BP ORFs. First, our data show that BP specific-CD4+ T cells are associated with a large and previously unrecognized breadth of responses, including hundreds of targets. Notably, fifteen distinct non-aP vaccine antigens were associated with reactivity comparable to that of the aP vaccine antigens. Second, the overall pattern and magnitude of CD4+ T cell reactivity to aP and non-aP vaccine antigens was similar regardless of aP vs wP childhood vaccination history, suggesting that the profile of T cell reactivity in adults is not driven by vaccination, but rather is likely driven by subsequent asymptomatic or sub-clinical infections. Finally, while aP vaccine responses were Th1/Th2 polarized as a function of childhood vaccination, CD4+ T cell responses to non-aP BP antigens vaccine responses were not, suggesting that these antigens could be used to avoid the Th2 bias associated with aP vaccination. Overall, these findings enhance our understanding of human T cell responses against BP and suggest potential targets for designing next-generation pertussis vaccines.

## INTRODUCTION

*Bordetella pertussis* (BP), a highly contagious pathogen responsible for thousands of deaths per year worldwide, is the causative agent of the respiratory tract infection known as whooping cough (CDC, 2023). Before the introduction of BP vaccines, about 200,000 cases/year of whooping cough (pertussis) caused by BP infection were reported in the United States alone. Introduction of vaccination in 1950 dramatically decreased disease incidence. The original whole-cell pertussis (wP) vaccine, based on inactivated bacteria adjuvanted with alum was phased out in the US because of adverse side effects, and replaced by acellular pertussis (aP) vaccine, a mixture of several different BP proteins (filamentous hemagglutinin [FHA], fimbriae 2/3 [Fim2/3], pertactin [PRN], and inactivated pertussis toxin [PtTox]) with alum, and universally adopted in the US in 1996 (Klein, 2014).

A resurgence in clinical pertussis was observed over the recent decades, affecting infants and children with incomplete immunization schedules, but also full vaccinated teenagers and young adults in ages coinciding with the implementation of the aP vaccine (Esposito et al., 2019; Kuehn, 2021). The increased disease incidence may in part reflect the change in vaccine composition, with aP vaccines limited to 4/5 antigens, and reportedly associated with a shorter protection from symptomatic disease (Chen and He, 2017; Chit et al., 2018; Jenkinson, 1988; Klein et al., 2013). However, several studies show that BP antibody responses for both vaccination and natural infection can be long lasting (Chit et al., 2018; Domenech de Celles et al., 2019; Wearing and Rohani, 2009; Wirsing von Konig et al., 1995). Additional studies reported that after booster vaccination, B cell memory and antibody responses decay rapidly within the first years but remain detectable and stable above baseline levels up to 12 years (Chen and He, 2017; Damron et al., 2020; Domenech de Celles et al., 2019; McAlister et al., 2021). Interestingly, protection against infection persists even after antibody titers have decreased (Dalby et al., 2010; Heininger et al., 2004a; Le et al., 2004), suggesting that a cell-mediated immune component contributes to BP protective immunity.

Intrinsic differences in T cell responses to BP in individuals primed with aP vs. wP vaccines have been described and several studies point to differences in terms of polarization evoked by infection or vaccination (Bancroft et al., 2016; da Silva Antunes et al., 2018; Higgs et al., 2012; Saso et al., 2021; van der Lee et al., 2018; Warfel et al., 2014; Wilk et al., 2019). Specifically, the two vaccines induce different qualities of T cell responses with aP-priming in infancy favoring a Th2 phenotype, in contrast to the Th1/Th17 phenotype associated with wP-priming vaccination (da Silva Antunes et al., 2018; da Silva Antunes et al., 2021a). Differences in polarization are maintained even in adulthood after repeated boosters (Ausiello et al., 2019; da Silva Antunes et al., 2018; Diavatopoulos and Edwards, 2017; Kapil and Merkel, 2019; Plotkin, 2018) possibly linked with immune imprinting (Consortium, 2019; da Silva Antunes et al., 2021b; Kapil and Merkel, 2019; Tian et al., 2018; Wilk et al., 2019).

Studies in animal models established that differential polarization, mucosal immunity and functionality of aP-induced adaptive responses are associated with decreased efficacy of aP vaccination in preventing infection and sub-clinical colonization, while retaining efficacy in the protection against pertussis disease (Solans and Locht, 2018; Warfel et al., 2014; Wilk et al., 2019; Zeddeman et al., 2020). Conversely, Th1/Th17 responses which are more prevalent in wP vaccination, appear to play a role in protection from both infection and disease (Ross et al., 2013). This is in contrast to Th2 responses that, while protecting from disease, have limited efficacy in diminishing colonization and bacterial carriage (Saso et al., 2021; Warfel et al., 2014; Wilk et al., 2019).

Pertussis is also common in older adults and particularly in patients with moderate/severe chronic obstructive pulmonary disease but frequently underdiagnosed and underreported (Kandeil et al., 2019; Macina and Evans, 2021; Wilkinson et al., 2021). Asymptomatic BP infections are highly prevalent in human populations (de Melker et al., 2006; Heininger et al., 2004b; Naeini et al., 2015; Palazzo et al., 2016; van Schuppen et al., 2022; Zhang et al., 2014), compatible with widespread sub-clinical reinfection. Adaptive immune responses associated with reinfections have been studied in the baboon and mouse models (Gregg et al., 2022; Warfel et al., 2014; Wilk et al., 2019; Zeddeman et al., 2020), but human data is scarce. It would be reasonable to expect that if asymptomatic human reinfections are relatively frequent regardless of previous vaccination status, the breadth of CD4+ T cell responses observed would be directed against aP and non-aP vaccine antigens alike.

Here we sought to fine map the landscape of human CD4+ T cell epitope and antigen recognition against BP, and ascertain if differences exist in the breadth, magnitude, and Th1/Th2 polarization between individuals originally primed with aP and wP vaccines. Definition of the targets of human T cell responses might also suggest potential antigens candidate for inclusion in novel vaccines.

## RESULTS

### Experimental design for a genome-wide screen of *Bordetella pertussis* human T cell epitopes

Previous studies characterized human CD4+ T cell reactivity to the four main antigens contained in the acellular pertussis vaccine, but little to no information is available regarding responses to other BP antigens. Here, we defined human CD4+ T cell reactivity spanning the entire BP proteome, by an approach previously used to draw a genome-wide map of human CD4+ T cell responses to *Mycobacterium tuberculosis* (MTB) (Lindestam Arlehamn et al., 2013).

The approach is based on predicting potential dominant CD4+ T cell epitopes from each ORFs encoded in the bacterial genome, based on their predicted promiscuous binding to HLA class II molecules (Paul et al., 2015). Previous studies demonstrated that this approach identifies the most dominant and prevalent epitopes, corresponding to approximately 50% of the total overall response (Grifoni et al., 2020; Oseroff et al., 2010). Accordingly, we synthetized a library encompassing a total of 24,877 peptides derived from 3,305 ORFs. The library was arranged in 133 pools of 188 15-mer peptides (hereafter called MegaPools; MP). Each MP was further divided in 8 pools of 22-24 individual peptides (hereafter called MesoPools; MS). A summary of the screening strategy is shown in **Supplementary Figure 1**.

The library was screened for CD4+ T cell reactivity utilizing peripheral blood mononuclear cells (PBMCs) collected in the 2013 to 2021 period from 40 participants, 21 males and 19 females, of 18 to 40 years of age. Based on clinical records and year of birth, 20 of the participants were immunized in childhood with a whole-cell pertussis (wP) vaccine, and 20 were originally immunized in childhood with an acellular pertussis (aP) vaccine. Demographics are summarized in **Supplementary Table 1a**.

### Large breadth of BP-specific CD4+ T cell responses in humans

CD4+ T cell reactivity was assayed directly *ex vivo* using an Activation Induced Marker (AIM) assay (**Supplementary Figure 1**), utilizing the combination of markers OX40+CD25+ (Dan et al., 2016), previously validated for epitope identification in the context of BP (da Silva Antunes et al., 2020). As done previously the threshold of positivity (TP) was based on the median twofold standard deviation of T cell reactivity in negative controls, corresponding to 285 cells per million of CD4+ T cells (0.0285%), and values above TP and with a stimulation index (S.I.)>2 were considered positive as previously described (da Silva Antunes et al., 2020; Tarke et al., 2022). An example of screening of the whole genome-wide library in a representative donor is shown in **Figure 1**. In this particular donor, we identified 32 positive MPs (**Figure 1A**). **Figure 1B** illustrates the deconvolution of one representative MP (MP#39), which yielded 3 positive MS. **Figure 1C** illustrates the deconvolution of the MS with the highest reactivity (MS#39.7), which identified 7 individual epitopes above the significance threshold. Overall, this particular donor recognized 148 different epitopes. **Figure 1D** shows the position of each individual epitope identified across the aligned BP genome, using the Tohama I and D420 BP strains as reference.

**Figure 1.**
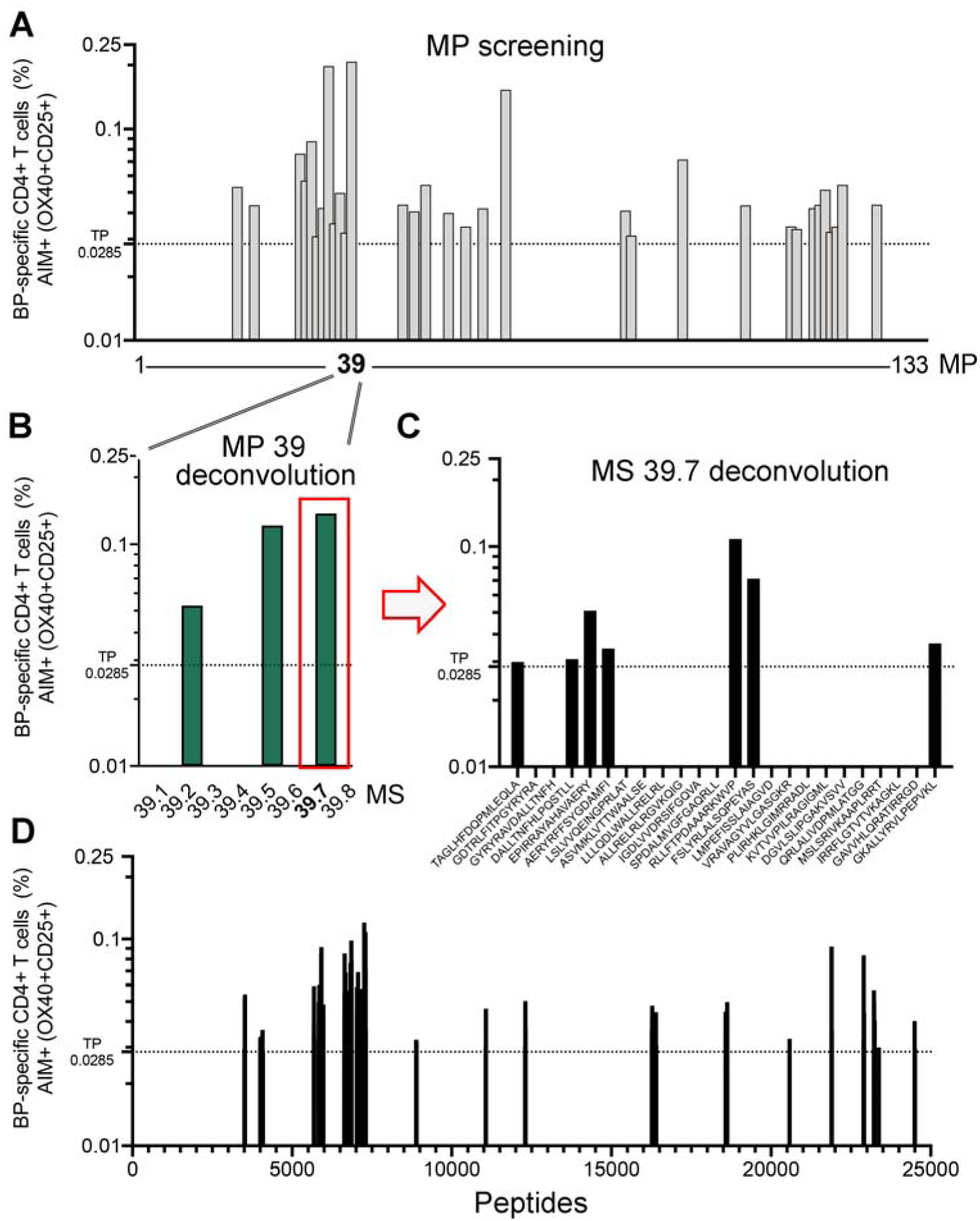
Schematics of BP whole genome-wide library screening. An example of the entire BP peptide library screening and epitope identification for a representative individual donor using AIM assay is shown. (A) Screening of entire library organized in 133 pools of 188 15-mer peptides (MegaPools; MP). (B) Deconvolution of one positive representative MP (MP#39) into 8 pools of 22-24 individual peptides (MesoPools; MS). (C) Deconvolution of one positive representative MS (MS#39.7) for assessment of individual peptide response (n=24). (D) Overall map of CD4+ T cell reactivity showing the position of each individual epitope identified across the aligned BP genome, using the Tohama I and D420 BP strains as reference. Associated percentage of response (magnitude) for each pool/peptide is indicated in y axis. Dotted lines represent the cut-off value associated with the threshold of positivity (TP).

To define the global pattern of immunodominance in the study cohort, we tested PBMCs from each donor with sets of the same peptide library and recorded the number of donors in which a positive response was detected. Peptides were defined as negative (not recognized), unconfirmed (recognized in only 1 donor), or confirmed positive (recognized in ≥2 donors). From hereafter, we took a conservative approach and focused on those epitopes that were reproducible (confirmed positive). The screening of the entire cohort revealed a total of 414 epitopes recognized in at least 2 donors, with 42 and 37 epitopes recognized in 3 or more than 4 donors, respectively (**Figure 2A** and **Supplementary Table 2**). Each individual epitope was mapped back to the individual BP ORFs of origin. A total of 239 ORFs were recognized. Of those 49 and 35 were associated with more than 2 and 3 epitopes respectively (**Figure 2B** and **Supplementary Table 3**). Parallel analyses quantified the total response across the entire cohort for each individual epitope or ORF. A total of 86 epitopes and 29 ORFs were required to account for 50% of the total response, 192 epitopes or 87 ORFs accounted for 75% of the response, and 301 epitopes or 152 ORFs were required to account for 90% of the total response (**Figure 2C, D**). Individual donors recognized, on average, 25 epitopes and 17 ORFs, underlining the large breadth of response to BP. Overall, the first quantitation of human CD4+ T cell responses to the whole BP genome revealed an unprecedented large breadth of antigens and epitopes recognized, especially when evaluating the diverse repertoire in individual donors.

**Figure 2.**
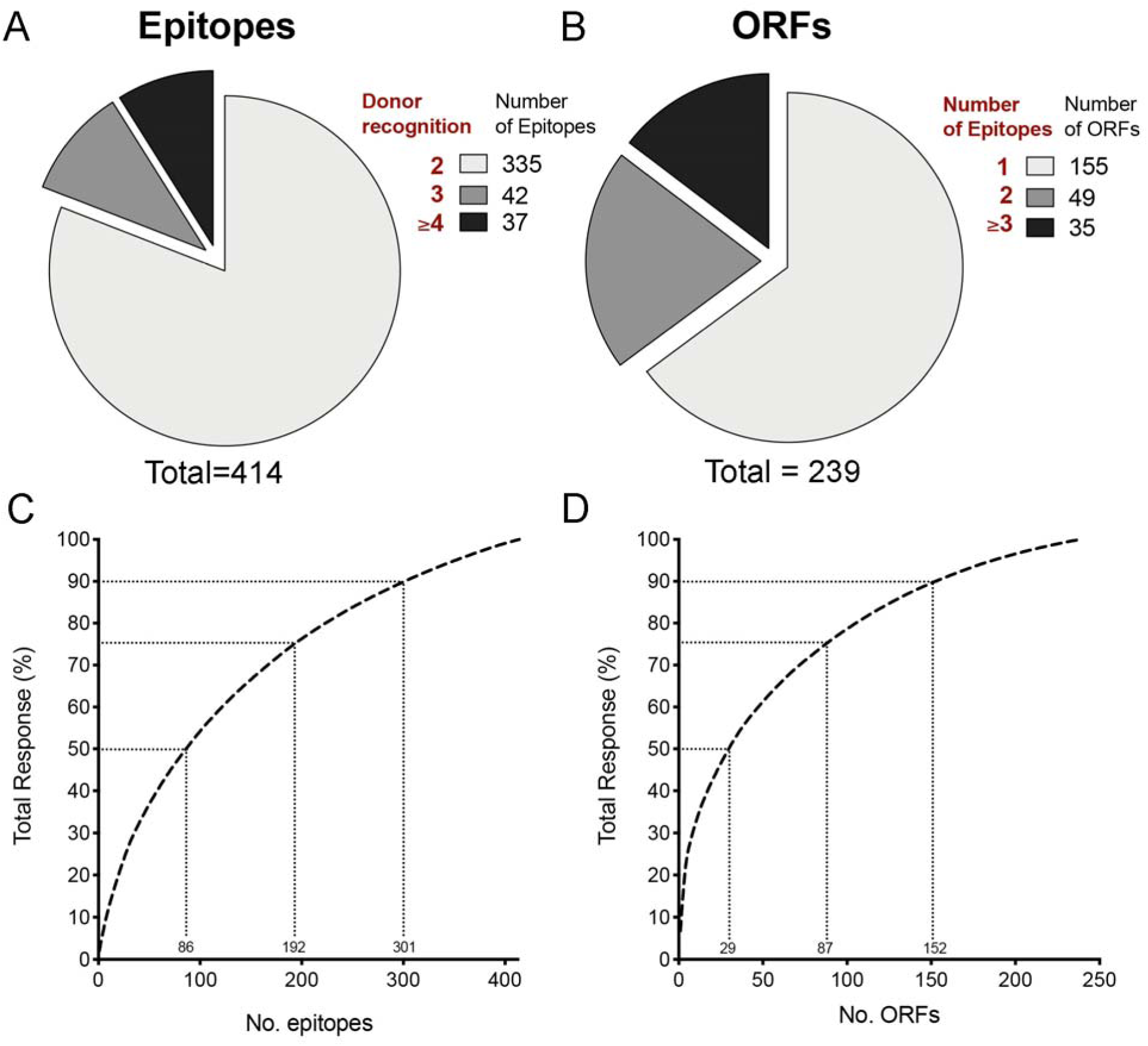
Large breadth of BP-specific CD4+ T cell responses in humans. (A) Dominance of epitope response across the entire cohort (n=40) indicated by proportion of donors who responded to the specified number of epitopes. (B) Proportion of the number of epitopes that account for the response of each individual ORFs (C) Breadth of epitope response ranked on the basis of % of total response (Black dashed line). Grey dotted lines indicate the top 50, 75 and 90 percent of total response and associated number of epitopes. (D) Breadth of antigen response ranked on the basis of % of total response (Black dashed line). Grey dotted lines indicate the top 50, 75 and 90 percent of total response and associated number of ORFs.

### Immunodominance in BP responses

Next, a linear map of the BP genome was used to visualize the combined reactivity of individual epitopes across all donors, showcasing both the overall magnitude of response and the specific region of each recognized ORF/antigen of origin (**Figure 3**). As expected, the known aP vaccine antigens (i.e., pertactin, PRN; two serotypes of fimbriae, Fim2/3; filamentous hemagglutinin, FHA; and pertussis toxin, PtTox) were amongst the most dominant antigens (**Figure 3**, in red and **Supplementary Table 3**). In particular, FHA was the antigen with the highest magnitude (6.76% of total response) and the most frequently recognized (42.5.0% of donors) of all the BP antigens. PRN and PtTox (ORF 1-5) were also associated with a high reactivity (4.70 % and 4.13% of total response, respectively) and high frequency of donor recognition (40.0% and 30%, respectively). Fim 3 and Fim 2 had the lowest reactivity (0.74% and 0.27% of total response, respectively) and were the least recognized among the aP vaccine antigens (10% and 5% of donors, respectively). Strikingly, the cumulative response of the 5 aP vaccine antigens only accounted for a minor fraction of the total CD4+ T cell response (16.6% of the total response; **Table 1**).

**Figure 3.**
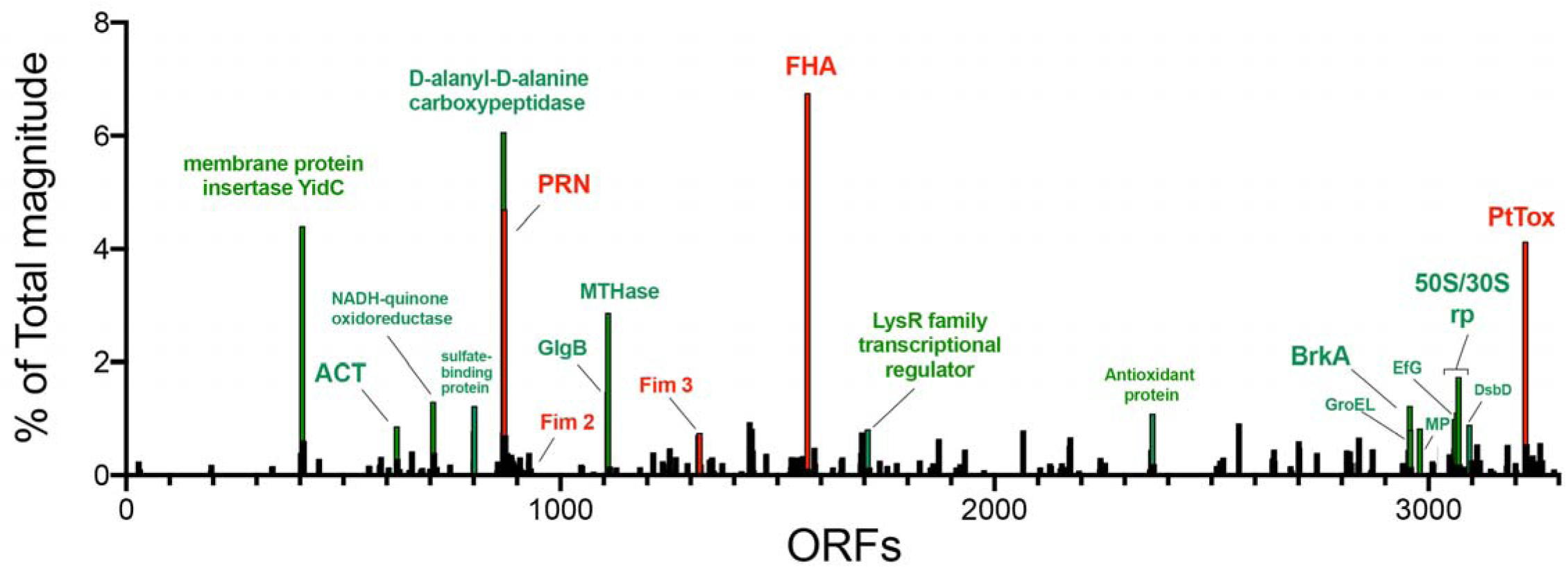
Immunodominance of BP specific-CD4+ T cell responses. Overall map of CD4+ T cell responses by antigen (ORF) reactivity across the entire cohort (n=40) showing the position of each individual ORF identified across the aligned BP genome, using the Tohama I and D420 BP strains as reference. Associated percentage of total response (all antigens recognized) for each ORF is indicated in y axis. Each bar represents an individual ORF and annotation of specific antigens is shown (red – aP vaccine antigens; green – dominant non-aP vaccine antigens).

**Table 1.**
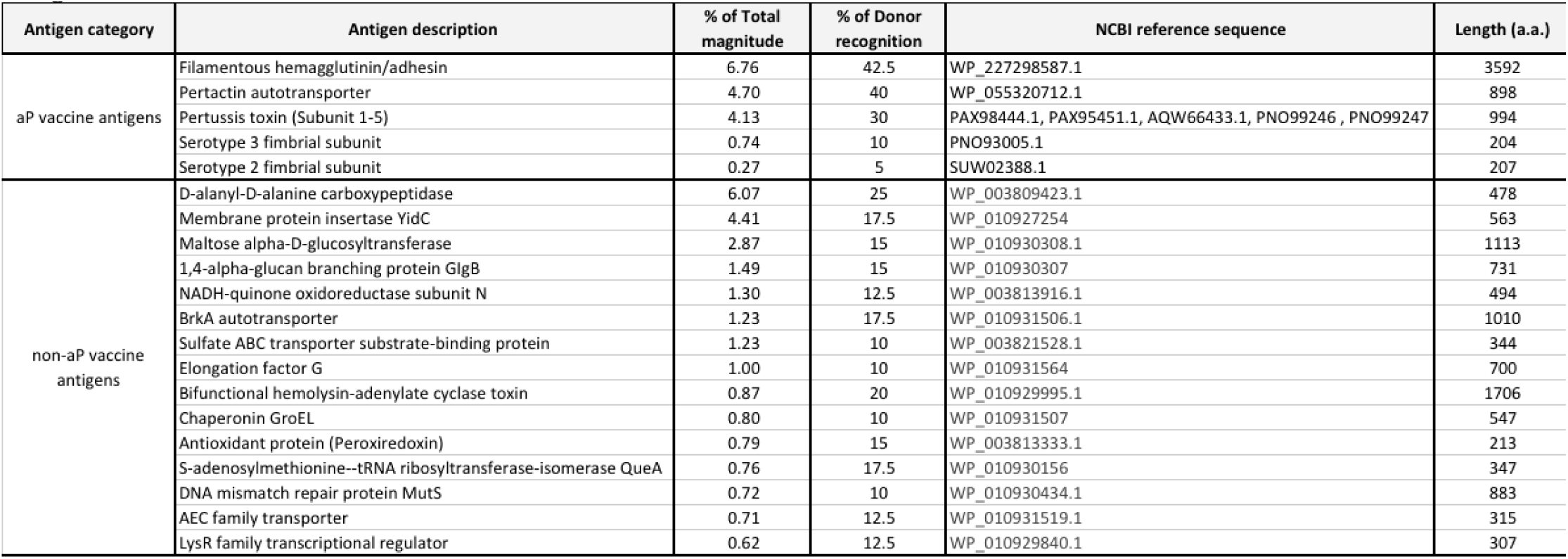
List of the immunodominant antigens with reactivity and frequency of recognition equal or better than aP vaccine antigens.

| Antigen category | Antigen description | % of Total magnitude | % of Donor recognition | NCBI reference sequence | Length (a.a.) |
| --- | --- | --- | --- | --- | --- |
| aP vaccine antigens | Filamentous hemagglutinin/adhesin | 6.76 | 42.5 | WP_227298587.1 | 3592 |
|  | Pertactin autotransporter | 4.70 | 40 | WP_055320712.1 | 898 |
|  | Pertussis toxin (Subunit 1-5) | 4.13 | 30 | PAX98444.1, PAX95451.1, AQW66433.1, PNO99246, PNO99247 | 994 |
|  | Serotype 3 fimbrial subunit | 0.74 | 10 | PNO93005.1 | 204 |
|  | Serotype 2 fimbrial subunit | 0.27 | 5 | SUW02388.1 | 207 |
| non-aP vaccine antigens | D-alanyl-D-alanine carboxypeptidase | 6.07 | 25 | WP_003809423.1 | 478 |
|  | Membrane protein insertase YidC | 4.41 | 17.5 | WP_010927254 | 563 |
|  | Maltose alpha-D-glucosyltransferase | 2.87 | 15 | WP_010930308.1 | 1113 |
|  | 1,4-alpha-glucan branching protein GlgB | 1.49 | 15 | WP_010930307 | 731 |
|  | NADH-quinone oxidoreductase subunit N | 1.30 | 12.5 | WP_003813916.1 | 494 |
|  | BrkA autotransporter | 1.23 | 17.5 | WP_010931506.1 | 1010 |
|  | Sulfate ABC transporter substrate-binding protein | 1.23 | 10 | WP_003821528.1 | 344 |
|  | Elongation factor G | 1.00 | 10 | WP_010931564 | 700 |
|  | Bifunctional hemolysin-adenylate cyclase toxin | 0.87 | 20 | WP_010929995.1 | 1706 |
|  | Chaperonin GroEL | 0.80 | 10 | WP_010931507 | 547 |
|  | Antioxidant protein (Peroxiredoxin) | 0.79 | 15 | WP_003813333.1 | 213 |
|  | S-adenosylmethionine--tRNA ribosyltransferase-isomerase QueA | 0.76 | 17.5 | WP_010930156 | 347 |
|  | DNA mismatch repair protein MutS | 0.72 | 10 | WP_010930434.1 | 883 |
|  | AEC family transporter | 0.71 | 12.5 | WP_010931519.1 | 315 |
|  | LysR family transcriptional regulator | 0.62 | 12.5 | WP_010929840.1 | 307 |

Conversely a high and broad reactivity was associated with non-aP vaccine antigens (**Figure 3**, in green and **Supplementary Table 3)**, and a total of 15 antigens were associated with reactivity and frequency of recognition similar or better than aP vaccine antigens (**Table 1**). Together, these 15 antigens accounted for about 25% of the total response, underlining the large heterogeneity of responses. ORFs associated with the highest magnitude were also associated with the highest frequency of responses (**Supplementary Figure 2**). The 15 antigens eliciting equivalent levels of responses as the vaccine antigens included ORFs from enzymes involved in roles such as cell wall and cell membrane assembly (D-alanyl-D-alanine carboxypeptidase and membrane protein insertase YidC, respectively) and carbohydrate metabolism (Maltose alpha-D-glucosyltransferase and 1,4-alpha-glucan branching protein GIgB). Among the dominant antigens were also adenylate cyclase toxin (ACT), BrkA, elongation factor G (EfG), and molecular chaperone GroEL which are associated with virulence and BP infection (DiVenere et al., 2022; Elder and Harvill, 2004; Moon et al., 2017). Overall, these results successfully re-identified aP vaccine antigens, and in addition greatly expanded the repertoire of antigens recognized by human BP-specific CD4 T cell responses.

### Similar recognition of aP and non-aP vaccine antigens as a function of priming vaccination in infancy

Half of our donor cohort was vaccinated in childhood with the wP vaccine, which is expected to generate responses targeting a wide range of BP antigens, while the other half was vaccinated in childhood with the aP vaccine, containing only four different antigens. To address whether the original priming would result in different repertoires of antigens recognized in adulthood, we compared responses from participants primed in infancy with aP versus wP vaccines for recognition of aP and non-aP vaccine antigens. No difference was detected for aP vaccine antigens (**Figures 4A-C**), considering either magnitude (sum of all reactivity of individual epitopes for a single donor) (**Figure 4A**), number of epitopes (**Figure 4B**) or ORFs recognized (**Figure 4C**). Similar magnitude, number of epitopes and ORFs between aP- and wP-primed donors were also observed when only responses to non-aP vaccine antigens were considered (**Figures 4D-F**). Overall, BP specific-CD4+ T cells responses did not differ as function of the original priming vaccination in infancy. The lack of significant differences between aP and wP originally primed donors, and the large breadth of responses, particularly in aP-primed donors is consistent with BP infection frequently occurring in vaccinated donors (de Melker et al., 2006; Naeini et al., 2015; Palazzo et al., 2016; van Schuppen et al., 2022).

**Figure 4.**
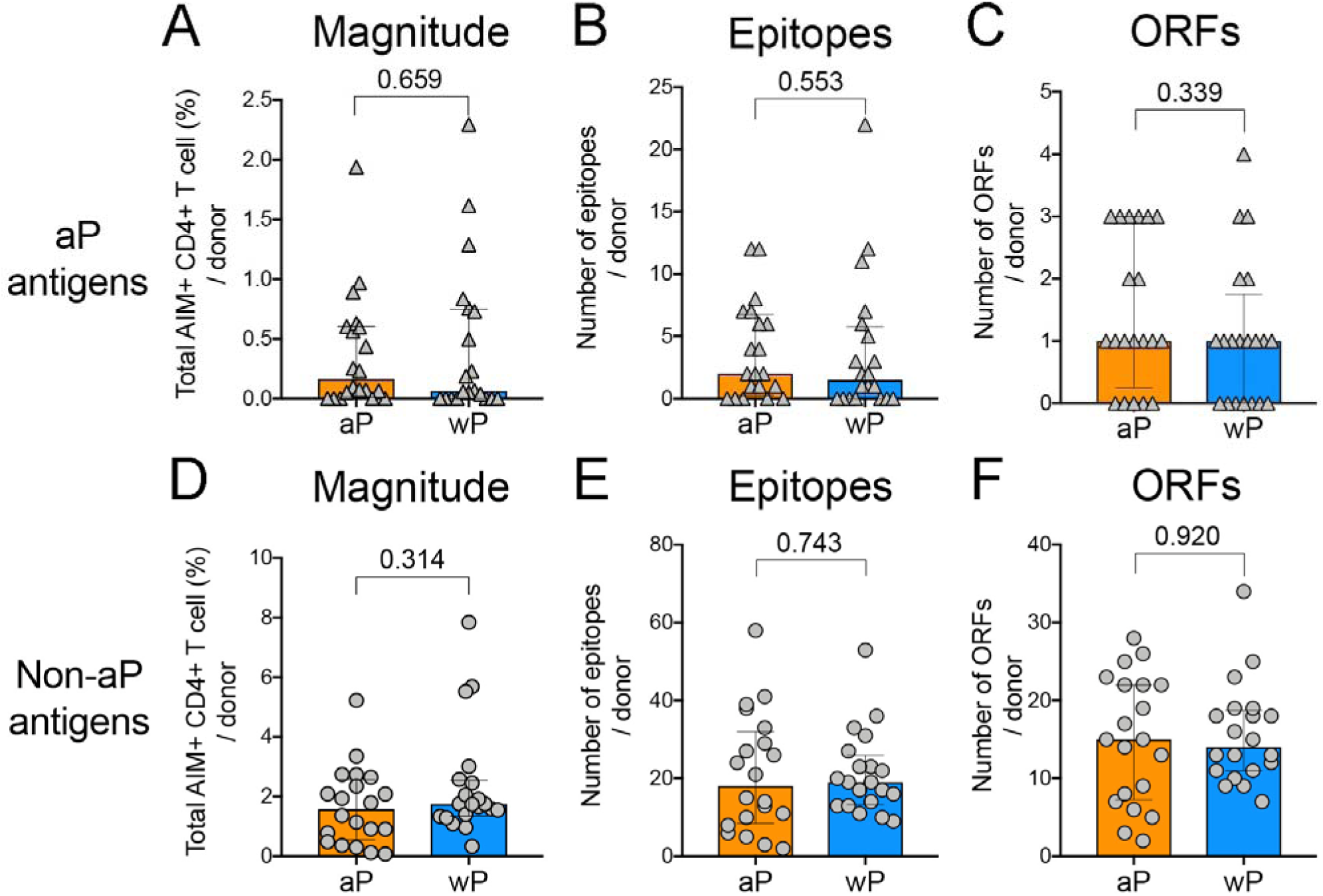
aP and non-aP vaccine antigens are similarly recognized in aP- and wP-primed donors. Graphs show comparison of responses between aP- and wP-primed donors in terms of (A,D) magnitude, (B,E) number of epitopes, and (C,F) number of ORFs for aP vaccine antigens (Grey triangles, upper panel) or non-aP vaccine antigens (Grey circles, bottom panel), respectively. Each symbol denotes an individual donor (n=40; 20 in each group). Bars represent geometric mean ± geometric SD. p values calculated by Mann-Whitney statistical analysis are indicated.

### Sequence conservation and immunogenicity of BP peptides

The degree of conservation of microbial sequences in different isolates and/or related species influence immunodominance, as conserved sequences have implicitly more opportunities to be recognized (Bui et al., 2007; Westernberg et al., 2016). Here, we performed a conservation analysis amongst different strains of BP or amongst different species of the genus *Bordetella*. Peptides were divided in subsets according to their immunogenicity or degree of conservation in different BP isolates or in related species. In terms of immunogenicity, peptides were categorized as previously described (i.e. negative (not recognized), unconfirmed (recognized in only 1 donor), or confirmed positive (recognized in ≥2 donors). In terms of conservation, peptides were classified as variable (<75% of homology), intermediate (75-95% of homology), or conserved (>95% of homology). Finally, peptides were further segregated as derived from aP vaccine antigens, or from non-aP vaccine antigens.

Twenty different BP strains with complete genomes and representative of the various clades of BP (Felice et al., 2022), and **Supplementary Table 4a**) were analyzed. Peptides from non-vaccine antigens are highly conserved (98.2-99.3% range) regardless of whether they are recognized by T cells or not (**Figure 5A)**. However, confirmed positive peptides are enriched in intermediate or conserved peptides (**Figure 5B)**, and non-reactive peptides are significantly enriched in variable peptides (**Figure 5C)**. Similar results were noted for peptides derived from aP vaccine antigens (**Figure 5D-F**), but the analysis significance is limited by the small number of peptides considered.

**Figure 5.**
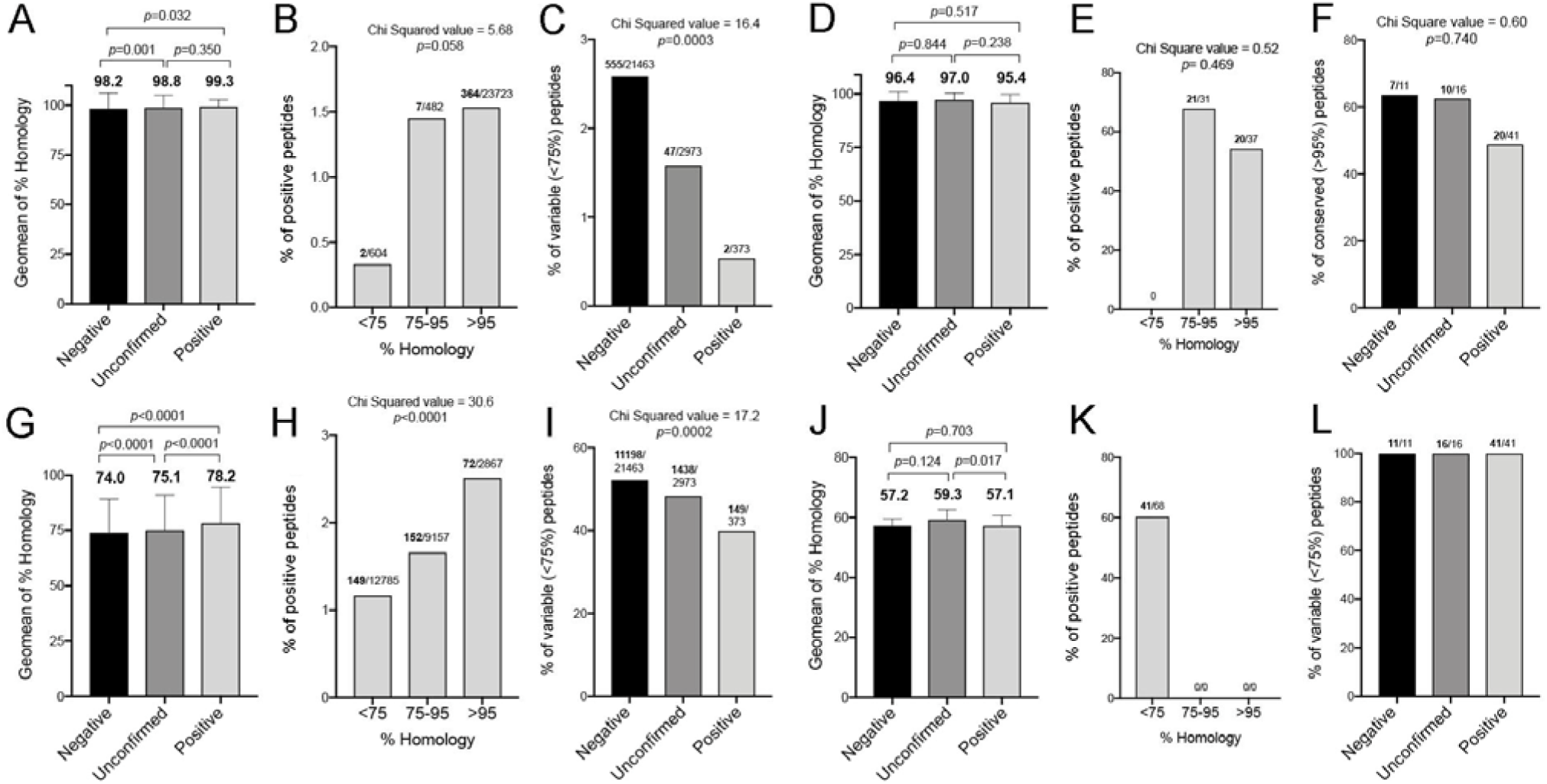
Sequence conservation is not a major driver of immunogenicity. Peptide homology amongst different BP strains or Bordetella genus was evaluated for the entire se of peptides tested in this study. (A) percent of homology across different BP strains for unrecognized (negative), unconfirmed or confirmed (positive) non-aP vaccine derived peptides (B) percent of positive peptides across different BP strains for peptide conservation in non-aP vaccine derived peptides. (C) percent of variable peptides across different BP strains for negative, unconfirmed or positive non-aP vaccine derived peptides. (D) percent of homology across different BP strains for negative, unconfirmed or positive aP vaccine derived peptides (E) percent of positive peptides across different BP strains for peptide conservation in aP vaccine derived peptides. (F) percent of conserved peptides across different BP strains for negative, unconfirmed or positive aP vaccine derived peptides. (G) percent of homology across different Bordetella for negative, unconfirmed or positive non-aP vaccine derived peptides (H) percent of positive peptides across different Bordetella for peptide conservation in non-aP vaccine derived peptides. (I) percent of variable peptides across different Bordetella for negative, unconfirmed or positive non-aP vaccine derived peptides. (J) percent of homology across different Bordetella for negative, unconfirmed or positive aP vaccine derived peptides (K) percent of positive peptides across different Bordetella for peptide conservation in aP vaccine derived peptides. (L) percent of variable peptides across different Bordetella for negative, unconfirmed or positive aP vaccine derived peptides. p values calculated by Kruskal-Wallis test adjusted with Dunn’s test for multiple comparisons are indicated.

We next investigated 22 different genomes (**Supplementary Table 4b)** of different species of the genus *Bordetella*. Peptides from non-vaccine antigens have low conserved sequences (74.0-78.2% range) regardless of whether they are recognized by T cells or not (**Figure 5G)**. As in the case of the strain analysis, confirmed positive peptides are enriched in conserved peptides, while non-reactive or unconfirmed peptides are enriched in variable peptides (**Figure 5H-I)**. Moreover, peptides from aP vaccine antigens are not conserved between species (<66.6%) with all peptides irrespectively of the T cell reactivity falling in the variable category (**Figure 5J-L)**. Overall, there is a significant level of sequence conservation within BP strains, but it is comparatively low among different species of *Bordetella*. While peptides that are linked with conserved sequences are more likely to be recognized, their contribution to the overall number of conserved peptides is minor (< 3%). Consequently, these peptides are not a major factor driving BP CD4+ T cell immunogenicity.

### Phenotypes associated with recognition of non-aP vaccine antigens

To characterize responses to the highest reactive non-aP vaccine antigens identified in this study, we generated a pool encompassing the top 170 epitopes which tested positive in at least 2 donors and hereafter denominated BP(E)R (See material and methods, and **Supplementary Table 5**). As control, we used a previously described MP (Bancroft et al., 2016; da Silva Antunes et al., 2018), containing epitopes exclusive from aP vaccine antigens [BP(E)VAC]. These MPs were tested in replicates of 3 independent experiments with PBMC from 20 subjects (10 aP and 10 wP) (**Supplementary Table 1b**). As expected, the BP(E)R pool yielded vigorous responses, which were not significantly different (*p*=0.11) from the BP(E)VAC pool in the AIM assay (**Figure 6A**), with 90 % and 75% of donor recognition, respectively.

**Figure 6.**
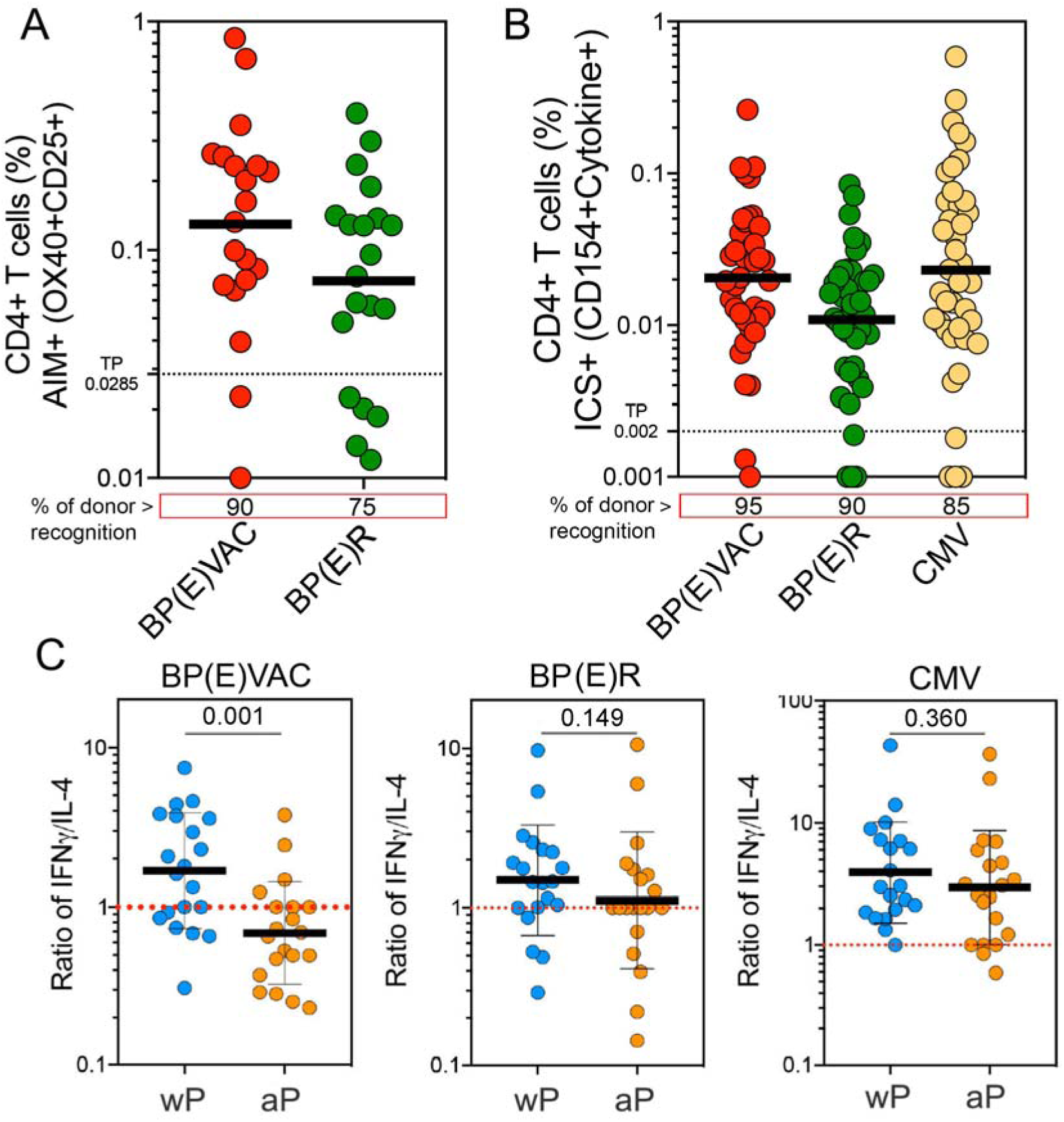
Th2 polarization is specific to the aP vaccine antigens, in individuals originally primed with aP vaccine. Antigen specific CD4+ T cell responses from aP and non-aP vaccine antigens were detected with BP(E)VAC and BP(E)R peptide pools respectively. (A) percentage of AIM+ (OX40+CD25+) CD4+ T cells after stimulation of PBMCs with peptide pools (n=20) (B) percentage of ICS+cytokine+ (CD154+) CD4+ T cells after stimulation of PBMCs with peptide pools (n=40). Black dotted lines represent the cut-off value associated with the threshold of positivity (TP) and percentage of donor recognition is indicated for each stimuli. (C) Polarization of CD4+ T cell responses represented as ratio of IFNγ/IL-4 cytokine response for each individual pool (*n*=40; 20 for aP and 20 for wP groups). Red dotted line indicates a Ratio=1. In all graphs, each circle represents a donor. Tick black lines represent geometric mean ± geometric SD, and p values were calculated by Mann-Whitney.

Previous studies highlighted differences in polarization patterns of responses to the aP vaccine antigens dependent on the original priming, with aP original vaccination being associated with a Th2 pattern and wP priming being associated with a Th1 profile (da Silva Antunes et al., 2018; da Silva Antunes et al., 2020). Here we addressed whether this difference in polarization was also noted with the non-aP vaccine antigens. Accordingly, CD4+ T cell responses to BP(E)VAC and BP(E)R were measured by intracellular cytokine staining (ICS) with phenotypic assessment of IFNγ, TNFα, IL-2 and IL-4 production among intracellular CD154+ (CD40L) cells in PBMC from an additional cohort of 40 subjects (20 aP and 20 wP) (**Supplementary Table 1c**). A MP against the ubiquitous pathogen cytomegalovirus (CMV) was used as an additional control of specificity.

As shown in **Figure 6B**, antigen-specific CD4+ T cell responses to BP(E)VAC and BP(E)R pools were also readily detected and similar in magnitude in terms of cytokine secretion with 95% and 90% of donor recognition, respectively. Interestingly, while responses to the aP vaccine antigens were Th1 polarized in wP-primed donors and Th2 polarized in aP-primed donors (**Figure 6C**), responses to the non-aP vaccine antigen pool BP(E)R were not polarized as function of priming vaccination in childhood, similar to CMV responses, which are known to be highly Th1 polarized (**Figure 6C**). Overall these results show that the Th2 polarization is specific to the aP vaccine antigens, in individuals originally primed with aP vaccine.

### CD4+ T cell responses against individual non-aP vaccine antigens

To further characterize responses to the most immunodominant non-aP vaccine antigens identified (**Table 1**), we focused on the antigens recognized in 5 or more donors (>12.5% donor recognition). Accordingly, we synthetized 11 MPs using sets of overlapping peptides encompassing the entire sequence of each individual antigen (ANT1-ANT11; 15-mer peptides overlapping by 10 residues; **Supplementary Table 6**). This unbiased approach allowed us to determine the overall reactivity of each antigen irrespective of the number of predicted peptides selected and tested from the initial screening library. In parallel, we used the above-mentioned BP(E)VAC peptide pool. These MPs were tested in replicates of 3 independent experiments with PBMC from 20 subjects (10 aP and 10 wP) using the AIM assay (**Supplementary Table 1d**).

To capture the overall response of each donor to all the 11 antigens, the magnitude of the individual MPs was combined (BP(O)ANT1-11). The results in **Figure 7A** show that responses to BP(O)ANT1-11 were robust and detected in all of donors, and yielded similar magnitude and donor recognition as BP(E)VAC. Among non-aP vaccine antigens, Maltose alpha-D-glucosyltransferase; MTHase (ANT3), BrkA (ANT6) and ACT (ANT7) were the most reactive antigens, with 35%, 50% and 70% of donors responding, respectively. These results could be in part attributed to the fact that in general, the most reactive antigens have the longest sequences (**Supplementary Figure 3**).

**Figure 7.**
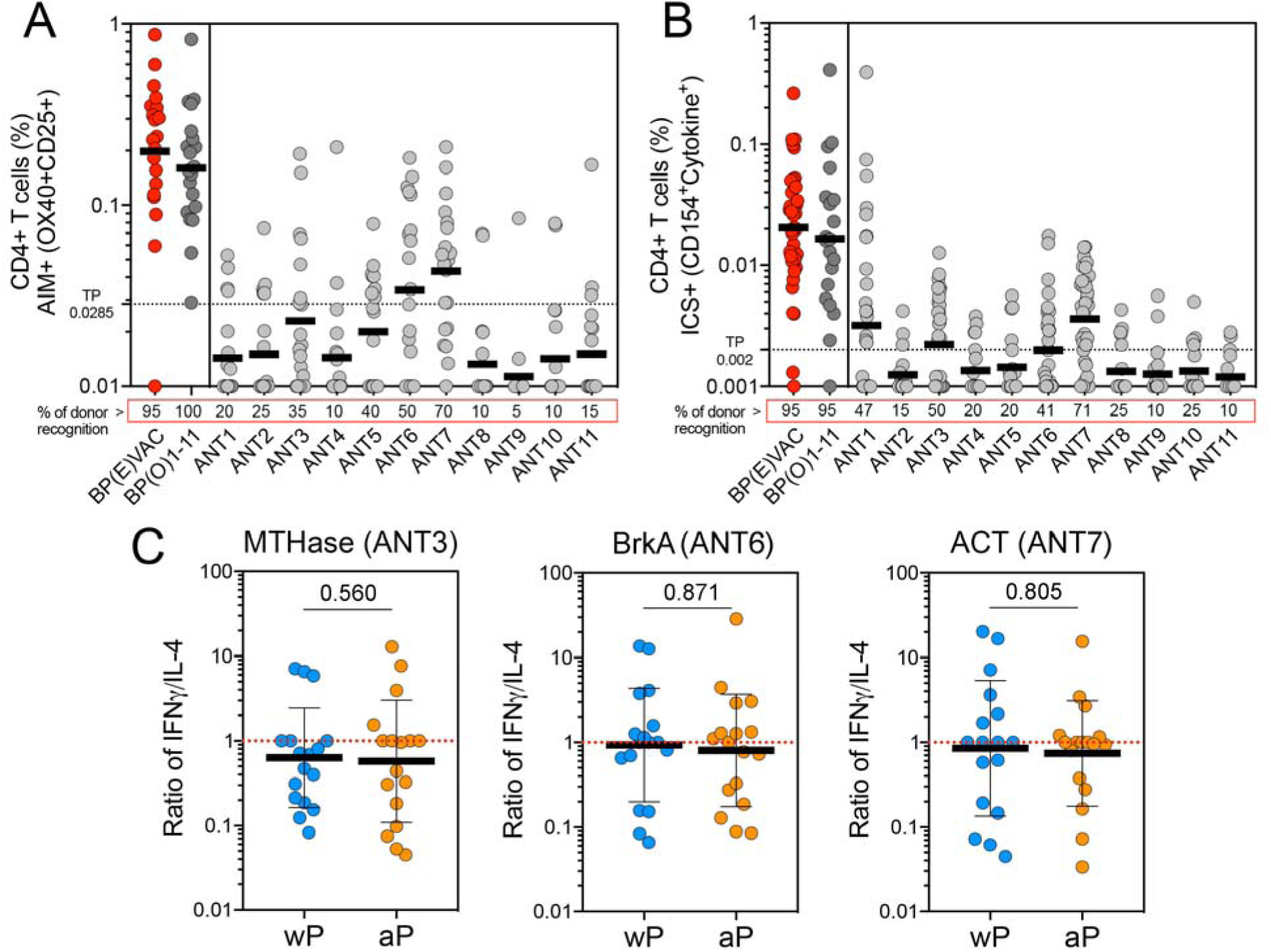
non-aP vaccine antigen responses are not polarized as function of priming childhood vaccination. Antigen specific CD4+ T cell responses from 11 individual non-aP vaccine dominant antigens were detected with overlapping (O) peptide pools and represented as the sum of all MPs responses (BP(O)1-15) or as each individual MP response (ANT1-ANT11). BP(E)VAC pool was used as control respectively. (A) percentage of AIM+ (OX40+CD25+) CD4+ T cells after stimulation of PBMCs with peptide pools (n=20) (B) percentage of ICS+cytokine+ (CD154+) CD4+ T cells after stimulation of PBMCs with peptide pools (BP(E)VAC, n=40; BP(O)1-15, n=20; ANT1, 3, 8, and 9, n=34; all other ANT, n=20). Black dotted lines represent the cut-off value associated with the threshold of positivity (TP) and percentage of donor recognition is indicated for each stimuli. (C) Polarization of CD4+ T cell responses represented as ratio of IFNγ/IL-4 cytokine response for each individual pool (*n*=34; 17 for aP and 17 for wP groups). Red dotted line indicates a Ratio=1. In all graphs, each circle represents a donor. Tick black lines represent geometric mean ± geometric SD, and p values were calculated by Mann-Whitney.

Detection of antigen-specific CD4+ T cell responses to the 15 non-aP vaccine antigens was also performed by ICS, in a subset of the previous cohort of 40 subjects (**Supplementary Table 1c**). Similar results to the AIM assay (**Figure 7A**) were observed for cytokine responses to all antigens combined (BP(O)ANT1-11), and to the 11 individual antigens (**Figure 7B**) with 50%, 41% and 71% of positive donor response for MTHase (ANT3), BrKA (ANT6) and ACT (ANT7), respectively. Consistent with the results shown above for the non-aP vaccine antigen pool BP(E)R, responses to non-aP vaccine individual antigens were not polarized as function of priming childhood vaccination (**Figure 7C**). These results confirm that responses to non-aP vaccine antigens are not Th2 polarized.

## DISCUSSION

Knowledge regarding human T cell responses to BP is largely limited to the four antigens contained in the aP vaccine, despite growing awareness of the importance of T cells in the control and prevention of symptomatic whooping cough disease (Chasaide and Mills, 2020; Fedele et al., 2015; Lambert et al., 2021; Solans and Locht, 2018; Warfel et al., 2014). Herein, we provide the first in-depth characterization of human CD4+ T cell responses to the whole BP proteome, spanning over 3,000 different ORFs.

In this study, we found that BP specific-CD4+ T cells in healthy young adults immunized with different pertussis vaccines are associated with a previously unrecognized and remarkable large breadth of responses. Tens of ORFs and hundreds of different peptides were recognized in the 40 donors studied. This remarkable large breadth of T cell responses parallels the large breadth of responses observed for another bacterial species (*Mycobacterium tuberculosis*: (Lindestam Arlehamn et al., 2013)), and it is a testament to the fact that T cells have the capacity to recognize most if not all foreign proteins.

Surprisingly, no differences were observed between wP- and aP-primed vaccinees in magnitude, breadth, or antigen repertoire of responses in both non-aP and aP vaccine antigens, indicating that the childhood priming does not drive the repertoire features detected in young adults. The broad responses to non-aP vaccine antigens in adults is consistent with the notion that repeated exposure and/or colonization might be a common occurrence throughout lifetime as suggested by the high prevalence of asymptomatic BP infections in human populations (de Melker et al., 2006; Heininger et al., 2004b; Naeini et al., 2015; Palazzo et al., 2016; van Schuppen et al., 2022; Zhang et al., 2014) and of adults being a reservoir for ongoing circulation of BP (Blanchard-Rohner, 2022). Lastly, this hypothesis is further supported by evidence from studies performed in baboon and mouse models that established that aP vaccination does not control infection, only symptomatic disease (Warfel et al., 2014; Wilk et al., 2019; Zeddeman et al., 2020). Since none of the donors participating in this study reported having experienced clinical whooping cough disease, the observed heterogeneity might reflect asymptomatic or sub-clinical infections.

While as expected, all the known aP vaccine antigens were re-identified, they did not account for the majority of the total CD4+ T cell response, and our studies identified fifteen different antigens that were recognized as vigorously as the aP vaccine antigens. These antigens included BrkA and ACT, known to play important roles in the pathogenesis of pertussis, as dominant targets of human T cell responses. Other antigens were associated with regulation of gene expression (LysR family transcriptional regulator and S-adenosylmethionine:tRNA ribosyltransferase) or enzymatic and catalytic activities related with peptidoglycan structure and remodeling (D-alanyl-D-alanine carboxypeptidase and membrane protein insertase YidC). Among other highly reactive targets we found several transporter proteins (sulfate ABC transporter substrate-binding protein and AEC family transporter) or antigens associated with Type I-IV secretion systems, a machinery system that translocate proteins and virulence factors (e.g. PtTox and ACT) from the cytoplasm to extracellular environment (Park et al., 2015). Indeed, transporter proteins and bacterial secretory systems appear to be frequently recognized by human T cell responses, possibly reflective of their prominent role in infection (Lindestam Arlehamn et al., 2013; Rolan and Tsolis, 2008; Saikh et al., 2006; Shepherd and McLaren, 2020).

What is the relation between the targets of CD4+ T cell and antibody recognition? The ACT antigen is also dominantly recognized by antibody responses in humans (Arciniega et al., 1991; Cherry et al., 2004), and additional CD4+ T cell antigens identified in this study are immunodominant for antibodies in mouse models (GroEL, EfG, and ribosomal proteins; (Raeven et al., 2015). In addition, a recent antigen discovery study using serum from convalescent baboons identified a total of 314 antigens targets of antibody responses, including several dominant targets of T cell responses in this study, such as GroEL, EfG, membrane protein insertase YidC, BrkA, and ACT. Because of the deterministic linkage between antibody and CD4+ T cell responses for large pathogens (Sette et al., 2008), our findings identify antigens that are targets of both antibody and T cell reactivity and should be considered in the design of upcoming pertussis vaccines.

The identification of new CD4+ T cell epitopes and immunodominant antigens not included in the aP vaccine, enabled us to develop tools to detect and characterize BP non-aP vaccine responses. While responses to the aP antigens were Th1 polarized in wP-primed donors and Th2 polarized in aP-primed donors as previously observed (Bancroft et al., 2016; da Silva Antunes et al., 2017; van der Lee et al., 2018), responses to non-aP antigens were not polarized as function of childhood priming vaccination. These results indicate that a Th2 phenotype is specific to aP vaccine antigens in individuals originally primed with aP vaccine. These results also suggest that the dominant non-aP vaccine antigens identified in this study could be used to induce a more balanced Th1/Th2 imprinting of memory BP responses even in subjects in which a dominant, long lasting Th2 response to the aP antigens is imprinted as a result of childhood aP vaccination.

Remarkably each individual tended to recognize a partially overlapping yet unique set of antigenic and epitope targets. The reasons for this donor-donor heterogeneity are not apparent, but might include past infection history, differences in HLA types, and influences of the microbiome on the repertoire of recognition. In this context, conservation amongst different *Bordetella* species is not a major driver of the dominantly recognized epitopes as in general, peptide homology or sequence conservation between different *Bordetella* isolates and BP was low, with only a small fraction of the highly conserved peptides associated with dominant responses. In addition, the majority of the *Bordetella* isolates studied are not typically associated with human infections, and therefore the high sequence homology is devoid of relevance.

We also anticipate that the newly established peptide pools will aid the research and characterization of CD4+ and CD8+ T cell responses in BP infection and controlled human BP infection/colonization initiatives in humans, and in the context of novel and/or whole-cell based vaccines candidates (Buddy Creech et al., 2022; Chasaide and Mills, 2020; Diks et al., 2023). Herein, we characterized CD4+ T cells responses directly ex vivo using high-throughput screening methodology, thus introducing minimal physiological perturbations and bias as a result of in vitro clonal expansion, and in combination with bioinformatic predictions of potential dominant epitopes binding HLA class II. This approach should be generally applicable to the study of T cell immune responses to other complex bacterial pathogens. Altogether, the fact that similar and broad repertoires are detected in aP-versus wP-originally primed subjects is compatible with the notion that asymptomatic infections occur similarly in the two groups. This appears to be independent of the reported difference between aP versus wP-originally primed subjects in terms of protection from symptomatic disease.

### Limitations of the study

A limitation of this investigation is the unknown history of previous BP exposure in the group of participants, including no direct evidence of re-infections. Our analysis is associated with samples limited to “steady-state” responses in adults and the monitoring and evolution of responses in a longitudinal fashion would be of considerable interest in future studies. Although these findings were validated with several different approaches, the epitope identification is limited to AIM-assessed peptide responses and serological analysis were not performed. Additional limitations of this study are the relatively small size and narrow age of the cohort investigated. The validation of the results in a cohort of wP primed individuals never boosted with aP vaccines, and a cohort clinically diagnosed with whooping cough would be desirable to generalize the findings.

## MATERIAL AND METHODS

### Study subjects

We recruited 62 healthy adults from the San Diego area (CA, USA) (**Supplementary Table 1**). All participants provided written informed consent for participation and relevant clinical medical history was collected and evaluated by the clinical coordinators through recording dates, type of pertussis vaccine, vaccination schedule, and questionnaires including if they ever experienced whooping cough disease. Individuals who had been diagnosed with BP infection at any given time in their life were excluded. All donors were originally vaccinated with either DTwP or DTaP in childhood and followed the recommended vaccination regimen (which is also necessary for enrollment in the California school system), which entails a Tdap booster immunization at 11-12 years and then every 10 years. In both groups, male and female subjects were included equally. PBMCs from each participant were isolated from whole blood or leukapheresis by density gradient centrifugation according to manufacturer instructions (Ficoll-Hypaque, Amersham Biosciences, Uppsala, Sweden) and cryopreserved for further analysis.

### Peptide prediction, synthesis, library assembly and pool preparation

Peptide selection was derived from either BP whole-genome predictions from the Tohama I strain (NCBI Accession number: NC_002929.2) or from a set of 256 unique open reading frames (ORF) from the recent clinical isolate D420 strain and not contained in Tohama I strain. These 256 unique ORFs have been identified in the lab of Dr. Tod Merkel (Gregg et al., 2022). BP genome-wide identification from Tohama I strain was performed by scanning for the presence of predicted HLA class II promiscuous binding peptides. MHC-peptide binding predictions were performed using publicly available tools hosted by the Immune Epitope Database (IEDB) Analysis Resource (Dhanda et al., 2019). Specifically, the prediction of peptides was established by the 7-allele HLA class II restricted method and by using peptides 15 residues in length and overlapping by 10 residues (Paul et al., 2015). Additional filtering using an epitope cluster analysis tool was performed to include unique peptides across all proteins with median percentile rank (cut-off of 10) predicted peptides for each antigen or at least 2 peptides per ORF (Dhanda et al., 2018). To remove redundant peptides, all peptides overlapping by 9 residues or more were placed into “variant clusters”. The most commonly occurring peptide was marked as the “representative” and the less-common peptides were marked as “variants”. Variants in each cluster were sorted by their alignment-start position and only synthesized once. Using these combined approaches, we selected and synthesized a total of 24,877 peptides, spanning 3,305 unique ORFs. The peptides were pooled and organized into a library of 1,064 MesoPools (MS) composed of 22-24 individual peptides, and also a library of 133 MegaPools (MP) composed of 8 MS. All individual peptides were synthesized by Mimotopes (Victoria, Australia) and resuspended to a final concentration of 1 mg/mL in DMSO.

### Whole genome screening study design

CD4+ T cell reactivity was assayed directly *ex vivo* using an Activation Induced Marker (AIM) assay previously validated for BP epitope discovery (da Silva Antunes et al., 2020). A summary of the screening strategy is shown in **Supplementary Figure 1**. PBMCs from each donor were tested with sets of the same peptide library After screening and identification of library pools that resulted in AIM+ reactive responses for each donor and based on cell availability we deconvoluted the top 30 MP and top 34 MS for each individual donor, which in preliminary analysis was shown to capture ≥75% and ≥90% of the total MP and MS library response, respectively. Overall, for each donor, an average of 764 peptides of the total library were tested and the position of each individual epitope identified mapped to the aligned BP genome using the Tohama I and D420 BP strains as reference. The total magnitude of response and localization of each recognized ORF/antigen across the entire cohort was performed by summing all the reactivity of individual epitopes across all donors.

### Generation of peptide pools for non-aP vaccine antigens

To validate and characterize responses to the most dominant non-aP vaccine epitopes identified in this study, we generated a pool encompassing 170 peptides [BP(E)R; E-experimentally defined; R-Rest of BP proteome] by selecting the top immunodominant epitopes recognized in at least 2 donors (**Supplementary Table 5**). To test the reactivity of 11 selected immunodominant non-aP vaccine antigens, we generated MPs of 15-mer peptides overlapping by 10 a.a. spanning the entire sequences of each individual antigen [BP(O)ANT1-11] (**Supplementary Table 6**). As a control, we also studied antigen-specific responses against a previously described MP (Bancroft et al., 2016; da Silva Antunes et al., 2018), containing epitopes exclusive from aP vaccine antigens [BP(E)VAC; E-experimentally defined; VAC-Vaccine antigens] and from the ubiquitous pathogen CMV (Yu et al., 2022). Individual peptides were synthesized by TC peptide lab (San Diego, CA).

### Activation Induced Marker (AIM) and Intracellular staining (ICS) assays

CD4+ T cell reactivity was assayed directly *ex vivo* using an Activation induced marker (AIM) assay utilizing the combination of markers OX40+CD25+ as previously described (Dan et al., 2016). This assay detects cells that are activated as a result of antigen-specific stimulation by staining antigen-experienced CD4+ T cells for TCR-dependent upregulation of OX40 and CD25 (AIM25) after an optimal time of 18–24 h of culture. Briefly, cryopreserved PBMCs were thawed, and 1 × 10^6^ cells/condition were immediately cultured together with peptide pools (2 μg/mL), individual peptides (10 μg/mL), or phytohemagglutinin-L (PHA) (10 μg/mL; Roche, San Diego, CA) and DMSO as positive and negative controls, respectively, in 5% human serum (Gemini Bio-Products) for 24 h. All samples were acquired on a ZE5 cell analyzer (Biorad laboratories, Hercules, CA) and analyzed with FlowJo software (Tree Star, Ashland, OR). AIM+ CD4+ T cells data were calculated as percentage of cells per million of CD4+ T cells. Background subtracted data were derived by subtracting the % of AIM+ cells percentage after each MP stimulation from the average of triplicate wells stimulated with DMSO. The Stimulation Index (SI) was calculated by dividing the % of AIM+ cells after peptide pool stimulation with the average % of AIM^+^ cells in the negative DMSO control. A positive response was defined as SI greater than 2 and AIM+ response above the threshold of positivity after background subtraction. The threshold of positivity (0.0285%) was calculated based on the median twofold standard deviation of T cell reactivity in negative DMSO controls according to previous published studies (da Silva Antunes et al., 2020; Tarke et al., 2022).

The intracellular cytokine staining (ICS) assay was performed as previously described (Tarke et al., 2022). PBMCs were cultured in the presence of antigen-specific MPs [1 mg/ml] in 96-well U-bottom plates at a concentration of 2×10^6^ PBMC per well. As a negative control, an equimolar amount of DMSO was used to stimulate the cells in triplicate wells and PHA (1mg/ml) stimulated cells were used as positive controls. After incubation for 24 hours at 37°C in 5% CO2, cells were incubated for additional 4 hours after adding Golgi-Plug containing brefeldin A, Golgi-Stop containing monensin (BD Biosciences, San Diego, CA) together with CD137 APC antibody (2:100; Biolegend, San Diego, CA). Cells were then stained on their surface for 30 min at 4°C in the dark, after that fixed with 1% of paraformaldehyde (Sigma-Aldrich, St. Louis, MO), permeabilized, and blocked for 15 minutes followed by intracellular staining for 30 min at room temperature. All samples were acquired on a ZE5 5-laser cell analyzer (Biorad laboratories, Hercules, CA) and analyzed with FlowJo software (Tree Star, Ashland, OR). Specifically, lymphocytes were gated, followed by single cells determination. T cells were gated for being positive to CD3 and negative for a Dump channel including in the same colors CD14, CD19 and Live/Dead staining. CD3+CD4+ were further gated based on a combination of each cytokine (IFNγ, TNFα, IL-2, and IL-4) with CD40L (CD154). The total cytokine response and T cell functionality was calculated from Boolean gating of single cytokines that was applied to CD3+CD4+ cells. The background was removed from the data by subtracting the average of the % of Cytokine+ cells plated in triplicate wells stimulated with DMSO. CD4+ T cell cytokine responses were background subtracted individually and found positive only if fulfilling the criteria of an SI greater than 2 and above a threshold of positivity (TP) of 0.002% for overall CD4+Cytokine+ cells. The TP for ICS was considered to be a positive response based on the median twofold standard deviation of T cell reactivity in negative DMSO controls.

The detailed information of all the antibodies used are summarized in **Supplementary Table 7,** and gating strategy is listed in **Supplementary Figure 4**. Gates were drawn relative to the negative and positive controls for each donor.

### Conservation analysis

The degree of conservation was performed amongst 20 strains representing different clades of BP or amongst 22 different species of the genus *Bordetella* (**Supplementary Table 4**). The protein sets for each BP strain were extracted using the NCBI’s assembly ID (https://www.ncbi.nlm.nih.gov/assembly) by retrieving FASTA files, and for each species were obtained by querying UniProt’s KnowledgeBase (UniProtKB - https://www.uniprot.org/uniprotkb) using each species’ taxon ID as the querying parameter. A bioinformatic analysis was conducted to ascertain the percent of homology for each individual peptide within all the different strains and species, using an IEDB tool PEPMatch (https://www.iedb.org; manuscript in preparation). Each peptide was searched to a sequence identity of <50% by allowing more than half of the sequence to have residue substitutions. Peptides were then divided in subsets according to their immunogenicity or degree of conservation in different isolates or related species. In terms of immunogenicity, peptides were arbitrarily divided in not recognized, subdominant (recognized in 1 donor), or dominant (recognized in >=2 donors). In terms of conservation, peptides were classified as variable (<75% of homology), intermediate (75-95% of homology), or conserved (>95% of homology). Finally, peptides were further segregated as derived from aP vaccine antigens, or from non-aP vaccine antigens. Results were plotted as geomean of percent homology or relative percent of the number of peptides in each subset.

### Statistical analysis

Comparisons between groups were performed using the nonparametric two-tailed, and unpaired Mann-Whitney test or Kruskal-Wallis test adjusted with Dunn’s test for multiple comparisons. Spearman’s rank correlation coefficient test was used for association analysis. Prism 8.0.1 (GraphPad, San Diego, CA, USA) was used for these calculations. Values pertaining to significance and correlation coefficient (R) are noted in the respective figure, and P < 0.05 defined as statistically significant.

### Study approval

This study was performed with approvals from the Institutional Review Board at La Jolla Institute for Immunology (protocols; VD-101-0513 and VD-059-0813). All participants provided written informed consent for participation and clinical medical history was collected and evaluated.

## AKNOWLEDGMENTS

Research reported in this publication was supported by the National Institute of Allergy and Infectious Diseases of the National Institutes of Health under Award Numbers U01 AI141995, U19 AI142742, and Contract Number 75N93019C00066. The content is solely the responsibility of the authors and does not necessarily represent the official views of the National Institutes of Health. We are grateful to all donors that participated in the study and the clinical studies group staff, particularly Gina Levi, as well as current and past members of the Peter and Sette laboratories for all their invaluable help.

## Supplementary Figures

**Supplementary Figure 1.**
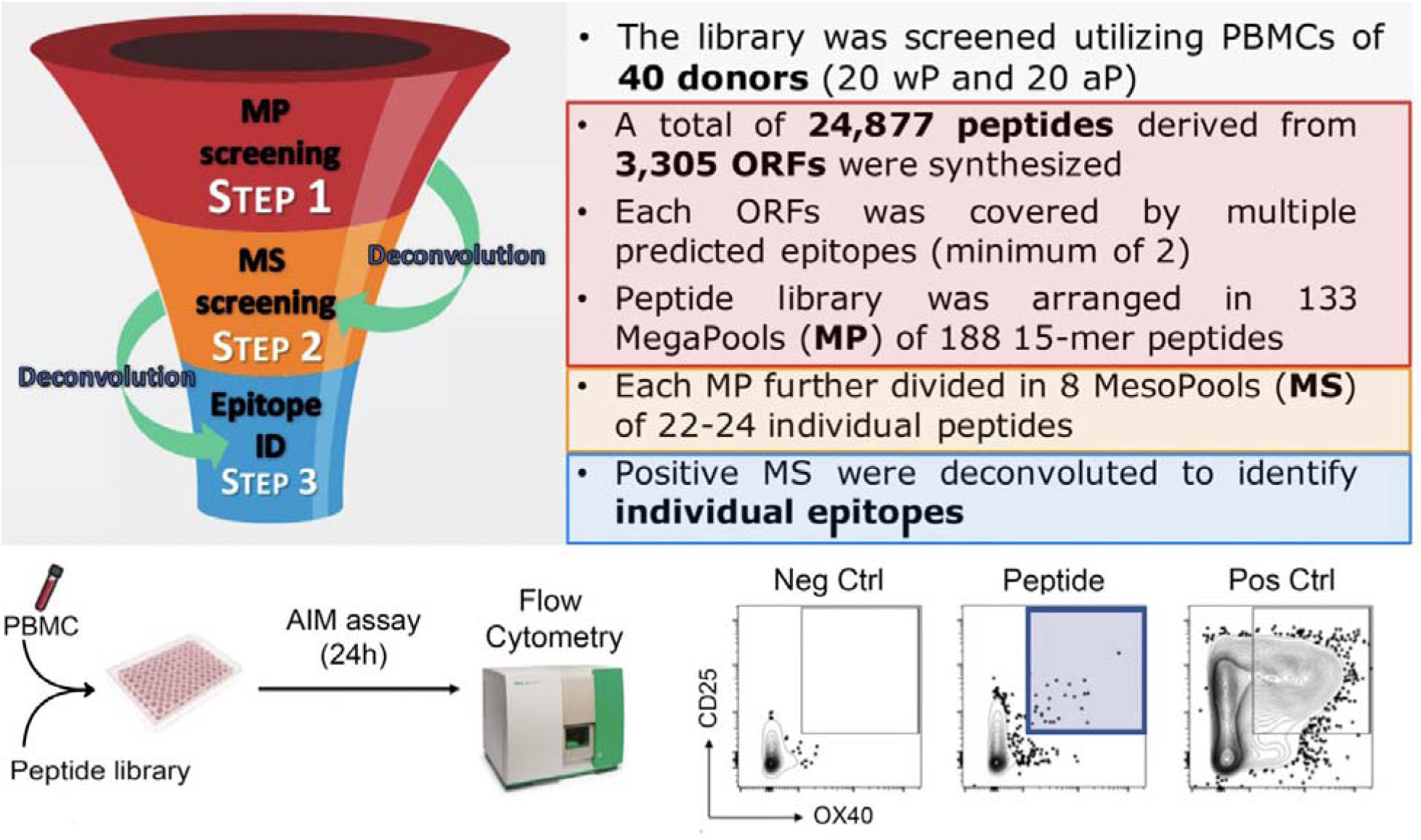
Schematic of BP genome-wide screening and summary of experimental design and strategy. CD4+ T cell reactivity spanning the entire BP proteome was assayed with a library of 24,877 peptides in 3 sequential steps, directly ex vivo using a high throughput Activation Induced Marker (AIM) assay flow cytometry methodology. Dot plots in the bottom right, show representative AIM+ CD4+ T cell responses after stimulation with a single peptide (red box) or with negative (DMSO) and positive (PHA) controls.

**Supplementary Figure 2.**
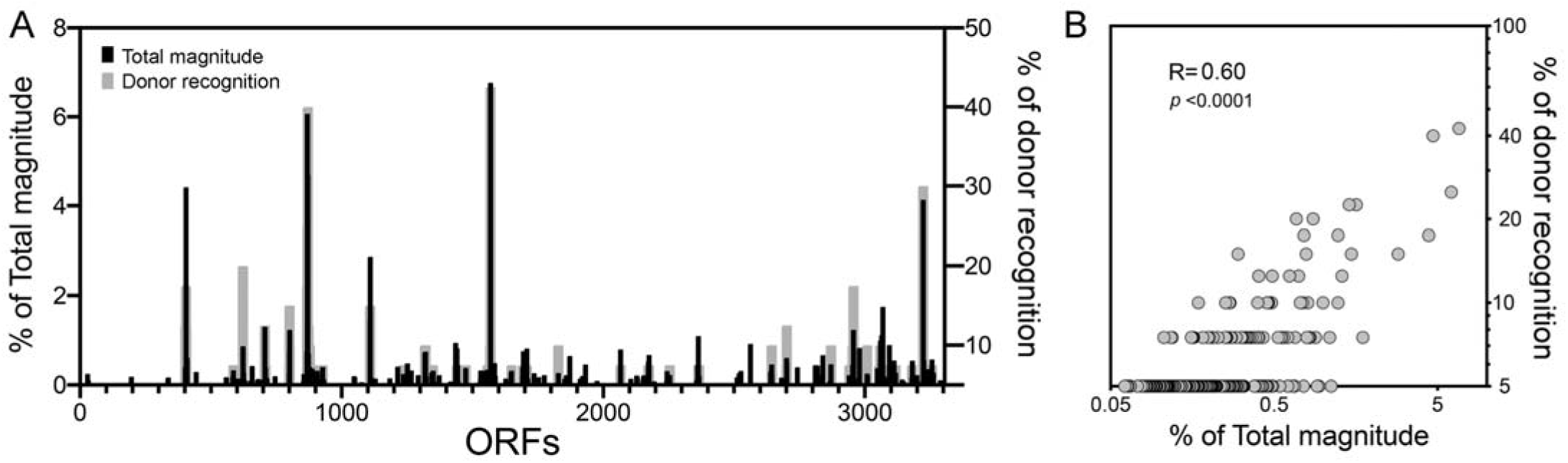
Immunodominance is associated with both magnitude and donor recognition. (A) Overall map of CD4+ T cell responses at antigen (ORF) level by percent of total magnitude (black bars, left axis) or percent of donor recognition (grey bars, right axis), across the entire cohort (n=40). Each bar represents an individual ORF identified across the aligned BP genome, using the Tohama I and D420 BP strains as reference. (B) Graph shows correlation between percentages of total magnitude and donor recognition. Each circle represents an individual ORF. R and p value expresses Spearman’s rank correlation coefficient test.

**Supplementary Figure 3.**
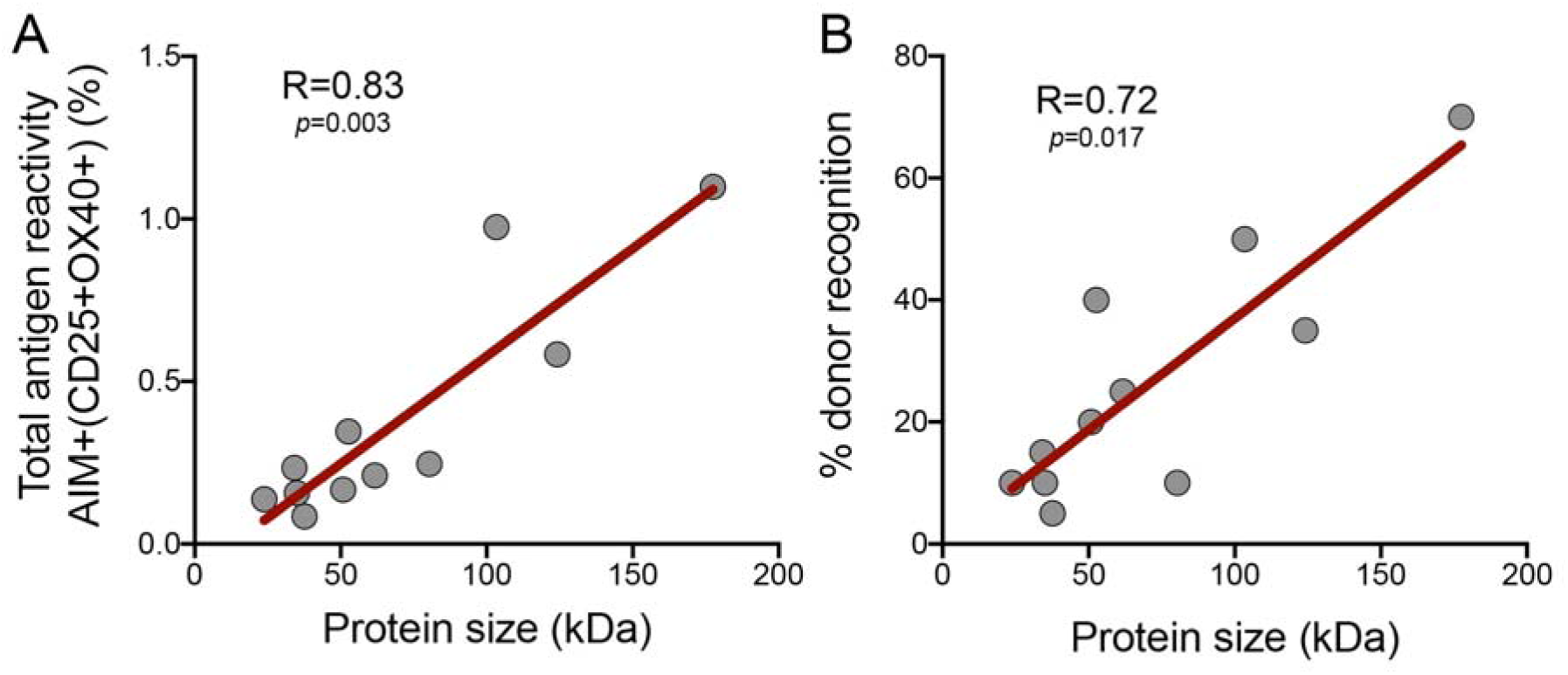
Immunodominant most reactive antigens have the longest sequences. Graphs shows correlation between protein size (kDa) and (A) percent of total antigen reactivity or (B) percent of donor recognition. Each circle represents responses of each of the 11 immunodominant overlapping peptide pools tested by AIM assay across all donors (n=20). R and p values express Spearman’s rank correlation coefficient test and the best fit is represented by a linear regression line (red).

**Supplementary Figure 4.**
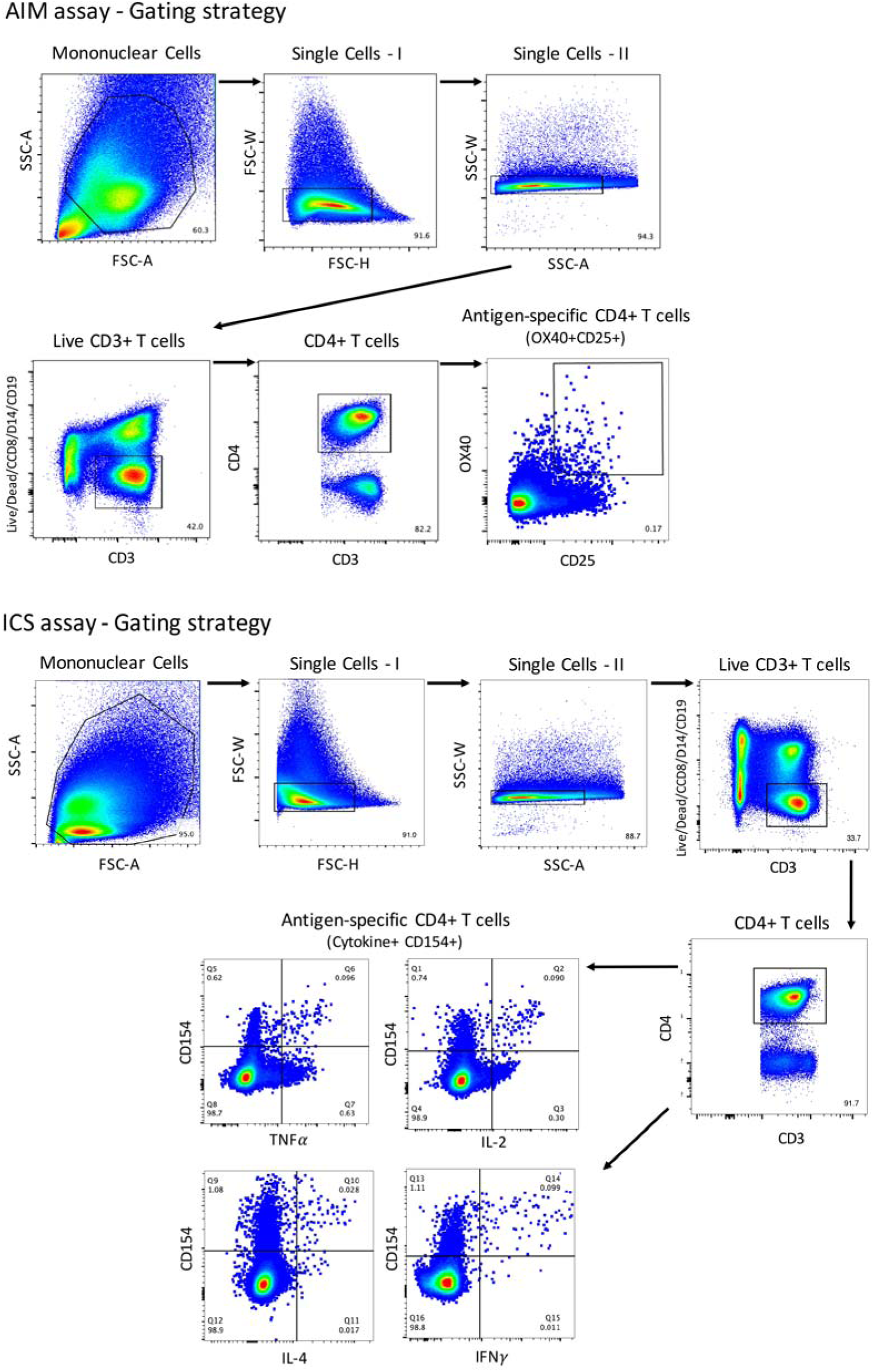
Illustrative flow cytometry gating strategy for the assessment of antigen-specific CD4+ T cell responses by AIM and ICS assays. Representative gating of reactive OX40+CD25+ and cytokine+(IFNγ, IL-2, TNFα and IL-4) CD154+ CD4+ T cells from donor PBMCs is shown. Briefly, for both AIM and ICS, mononuclear cells were gated out of all events followed by subsequent singlet gating. Live CD3+ cells were gated as Live/Dead-CD14-CD8-CD19-CD3+. Cells were then gated as CD4+CD3+. For AIM assay, antigen-specific cells were defined as OX40+CD25+ CD4+ T cells (AIM+) after antigen stimulation, and frequencies calculated as percent of total CD4+ T cells after background subtraction. For ICS assay, antigen-specific cytokine producing cells were defined as Cytokine+ and CD154+ CD4+ T cells after antigen stimulation, and frequencies calculated as percent of total CD4+ T cells after background subtraction.

